## Supplemental Material for "Structural and thermodynamic analyses of the β-to-α transformation in RfaH reveal principles of fold-switching proteins"

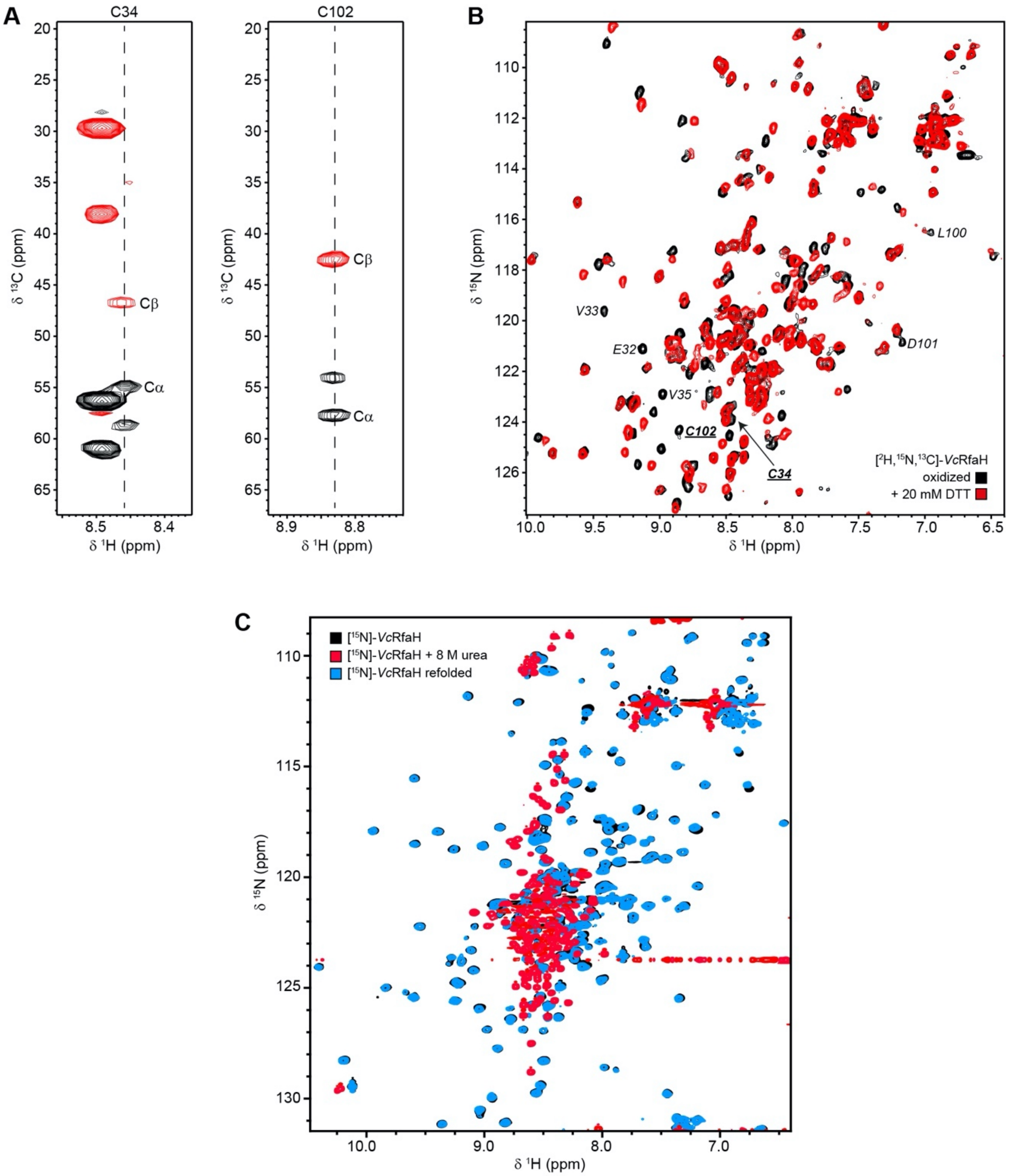


**Figure 1 – Figure supplement 1.** Disulfide bridge formation in *Vc*RfaH. (**A**) Strips of the HNCACB experiment corresponding to *Vc*RfaH residues C34 and C102, respectively. Signals arising from the cystein's Cα and Cβ carbons (indicative of a cysteine in a disulfide-bridge) are labeled. (**B**) [$^{1}$H, $^{15}$N]-HSQC spectra of [$^{2}$H, $^{13}$C, $^{15}$N]-*Vc*RfaH in the absence (black) or presence (red) of 20 mM DTT. Signals of the two disulfide bridge forming residues, C34 and C102, and their sequential neighbors are labeled. (**C**) Refolding of *Vc*RfaH under reducing conditions. [$^{1}$H, $^{15}$N]-HSQC spectra of 150 µM $^{15}$N-*Vc*RfaH (black), 39 µM $^{15}$N-*Vc*RfaH after incubation in the presence of 8 M urea for 24 hours (red), and 40 µM $^{15}$N-*Vc*RfaH upon refolding (cyan).

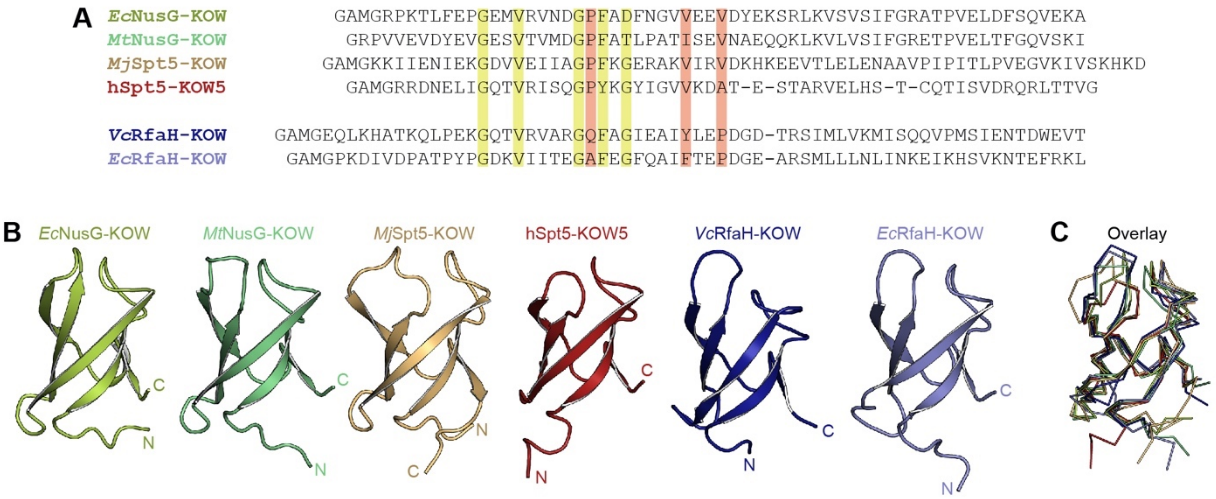


**Figure 1 – Figure supplement 2.** Structure comparison of KOW domains used in this study. (**A**) Structure-based sequence alignment of the KOW constructs used in this study. Highly conserved residues are highlighted in green, residues that differ among NusG-KOW/Spt5-KOW and RfaH-KOW domains are colored red. (**B**) Structures of the six KOW domains shown in cartoon representation. N- and C-termini are labeled. PDB-IDs: 2JVV (*Ec*NusG-KOW), 2MI6 (*Mt*NusG-KOW), 4ZN3 (*Mj*Spt5-KOW), 2E70 (hSpt5-KOW5), 2LCL (*Ec*RfaH-KOW), 6TF4 (*Vc*RfaH-KOW). (**C**) Structural alignment of the six KOW domains. The proteins are shown as ribbons. Orientation and colors as in (B).

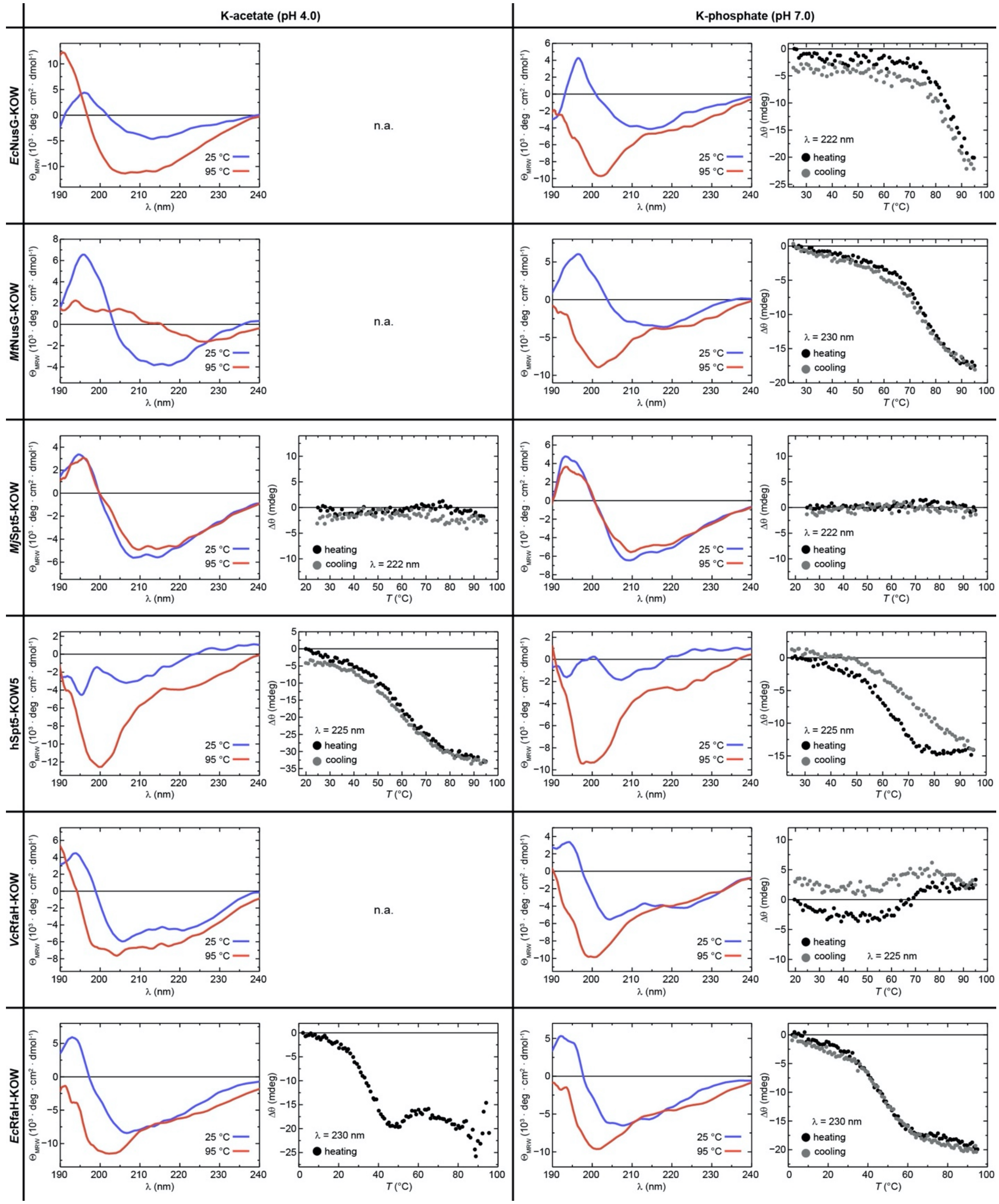


**Figure 2 – Figure supplement 1.** Reversibility of thermal unfolding. The graphs show CD spectra of the six KOW domains at 25 °C (blue) and at 95 °C (red) together with the change in ellipticity,

Δθ, during heating from 25 °C to 95 °C (filled black circles) and subsequent cooling to the initial temperature (filled grey circles), each at pH 4.0 (left) and pH 7.0 (right). When aggregation was already apparent from the CD spectra acquired at 95 °C (i.e. the shape of the spectrum did not correspond to that of an unfolded protein), no thermal unfolding/refolding curves were recorded (n.a.). Due to its hyperthermophilic source organism, *Mj*Spt5-KOW could not be denatured at either pH. The wavelength for monitoring the transition was chosen based on the largest difference between the spectra of the folded and unfolded state (for details see Materials & Methods).

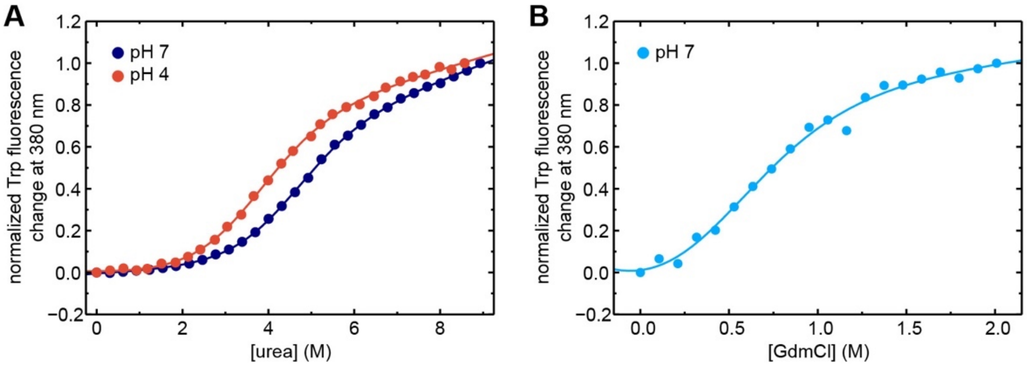


**Figure 3 – Figure supplement 1.** Chemical unfolding of *Vc*RfaH-KOW monitored by change in Trp fluorescence. (**A**), (**B**) The curves show the normalized Trp fluorescence change at 380 nm of *Vc*RfaH-KOW, obtained after over-night incubation of the protein in the presence of increasing concentrations of (A) urea at pH 4.0 (filled blue circles) or pH 7.0 (filled red circles) or (B) GdmCl at pH 7.0 (filled light blue circles). The lines represent fits to a two-state unfolding model.

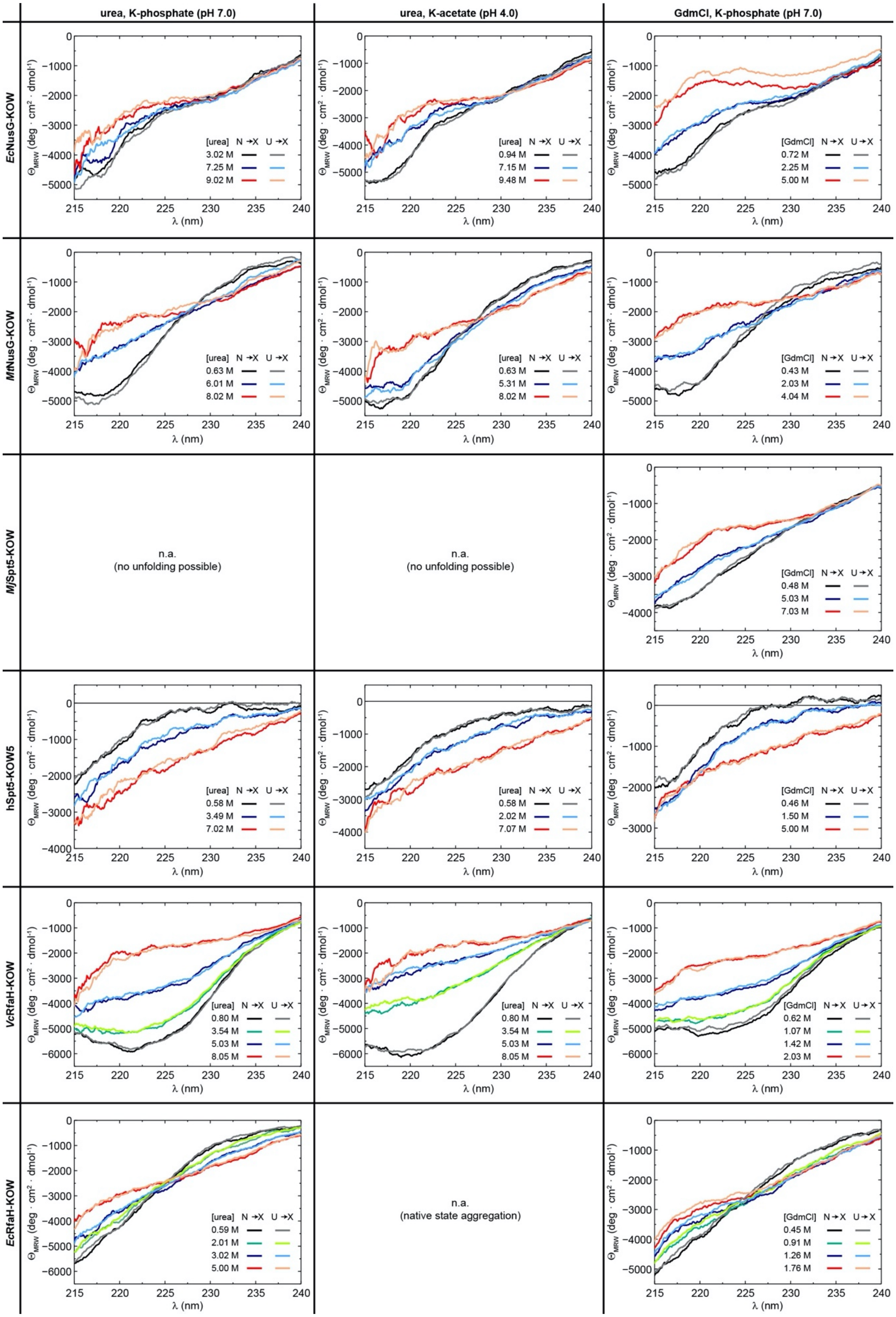

urea, K-phosphate (pH 7.0)
urea, K-acetate (pH 4.0)
GdmCl, K-phosphate (pH 7.0)
EcNusG-KOW
MfNusG-KOW
MjSpt5-KOW
hSpt5-KOW5
VcRfaH-KOW
EcRfaH-KOW
[urea] N →X U →X
3.02 M
7.25 M
9.02 M
0.94 M
7.15 M
9.48 M
[GdmCl] N →X U →X
0.72 M
2.25 M
5.00 M
0.63 M
6.01 M
8.02 M
0.63 M
5.31 M
8.02 M
0.43 M
2.03 M
4.04 M
n.a.
(no unfolding possible)
n.a.
(no unfolding possible)
0.48 M
5.03 M
7.03 M
0.58 M
3.49 M
7.02 M
0.58 M
2.02 M
7.07 M
0.46 M
1.50 M
5.00 M
0.80 M
3.54 M
5.03 M
8.05 M
0.80 M
3.54 M
5.03 M
8.05 M
0.62 M
1.07 M
1.42 M
2.03 M
0.59 M
2.01 M
3.02 M
5.00 M
n.a.
(native state aggregation)
0.45 M
0.91 M
1.26 M
1.76 M
Θ_MRW (deg · cm² · dmol⁻¹)
λ (nm)

**Figure 3 – Figure supplement 2.** Reversibility of chemical denaturation. CD spectra of the six protein domains acquired at the indicated denaturant concentration and buffer. In order to check the reversibility, two spectra at identical denaturant concentration were obtained by adding the native protein from a solution containing no denaturant to the desired denaturant concentration (N → X), or by adding the unfolded protein from a solution containing 10 M urea/8 M GdmCl to a solution containing the desired denaturant concentration (U → X). The color code is indicated.

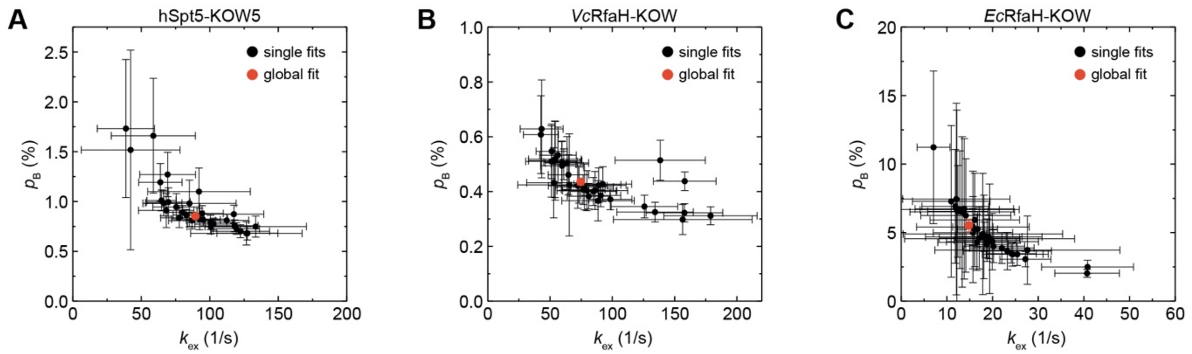


**Figure 5 – Figure supplement 1.** Extended CEST analysis of hSpt5-KOW5, *Vc*RfaH-KOW, or *Ec*RfaH-KOW. Plots of $k_{ex}$ vs. the population of the minor species ($p_B$) obtained from individual fits (black symbols) or a global fit (red symbol) of the CEST profiles of (**A**) hSpt5-KOW5, (**B**) *Vc*RfaH-KOW, and (**C**) *Ec*RfaH-KOW. Error bars represent the standard deviation of the fits.

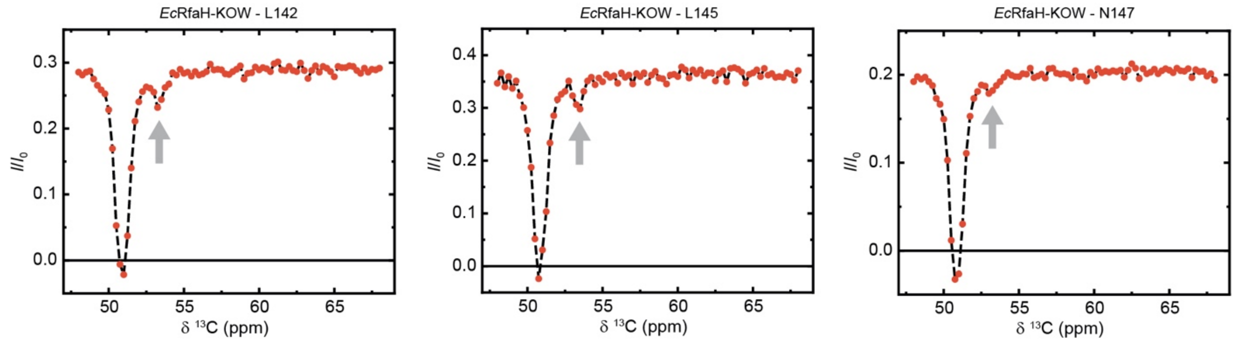


**Figure 6 – Figure supplement 1**. Exemplary traces of CEST experiments recorded on $^{13}$Cα carbons of $^{13}$C-*Ec*RfaH-KOW. The arrows indicate the positions of the minor species dips.

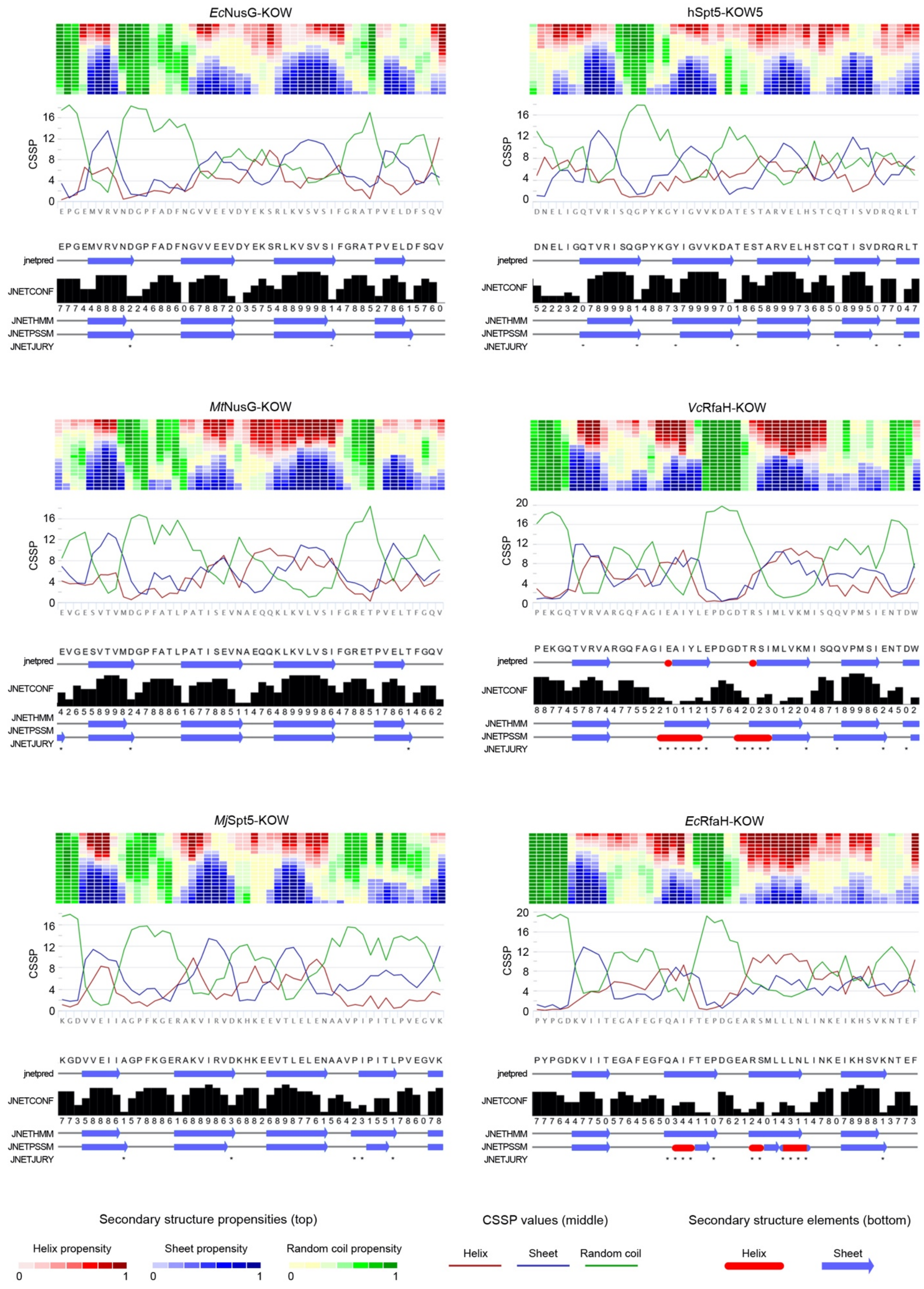
EcNusG-KOW
hSpt5-KOW5
MtNusG-KOW
VcRfaH-KOW
MjSpt5-KOW
EcRfaH-KOW
CSSP
jnetpred
JNETCONF
JNETHMM
JNETPSSM
JNETJURY
Secondary structure propensities (top)
CSSP values (middle)
Secondary structure elements (bottom)
Helix propensity
Sheet propensity
Random coil propensity
Helix
Sheet
Random coil

**Figure 6 – Figure supplement 2.** Secondary structure predictions for the six KOW domains used in this study. For each domain two predictions were caried out: (i) By Net-CSSP (Kim et al., 2009), top. The contact-dependent secondary structure propensity (CSSP) of each domain is plotted against die amino acid sequence (red: helices; blue: beta structures; green: random coil). The heat map above each graph displays the propensity of each amino acid to adopt helical (red), beta (blue), or random coil (green) structures using a gradient from dark (high propensity) to light (low propensity) colors. (ii) By Jpred 4 (Drozdetskiy et al., 2015), bottom. The predicted secondary structure elements are shown below the amino acid sequence (red: helices; blue: beta structures). JNetPRED: consensus prediction; JNetCONF: confidence estimate for the prediction (high values correspond to high confidence); JNetHMM: profile prediction based on hidden Markov model (HMM), JNetPSSM: profile prediction based on position-specific scoring matrix (PSSM); JNetJURY: an asterisk indicates significantly different primary predictions.

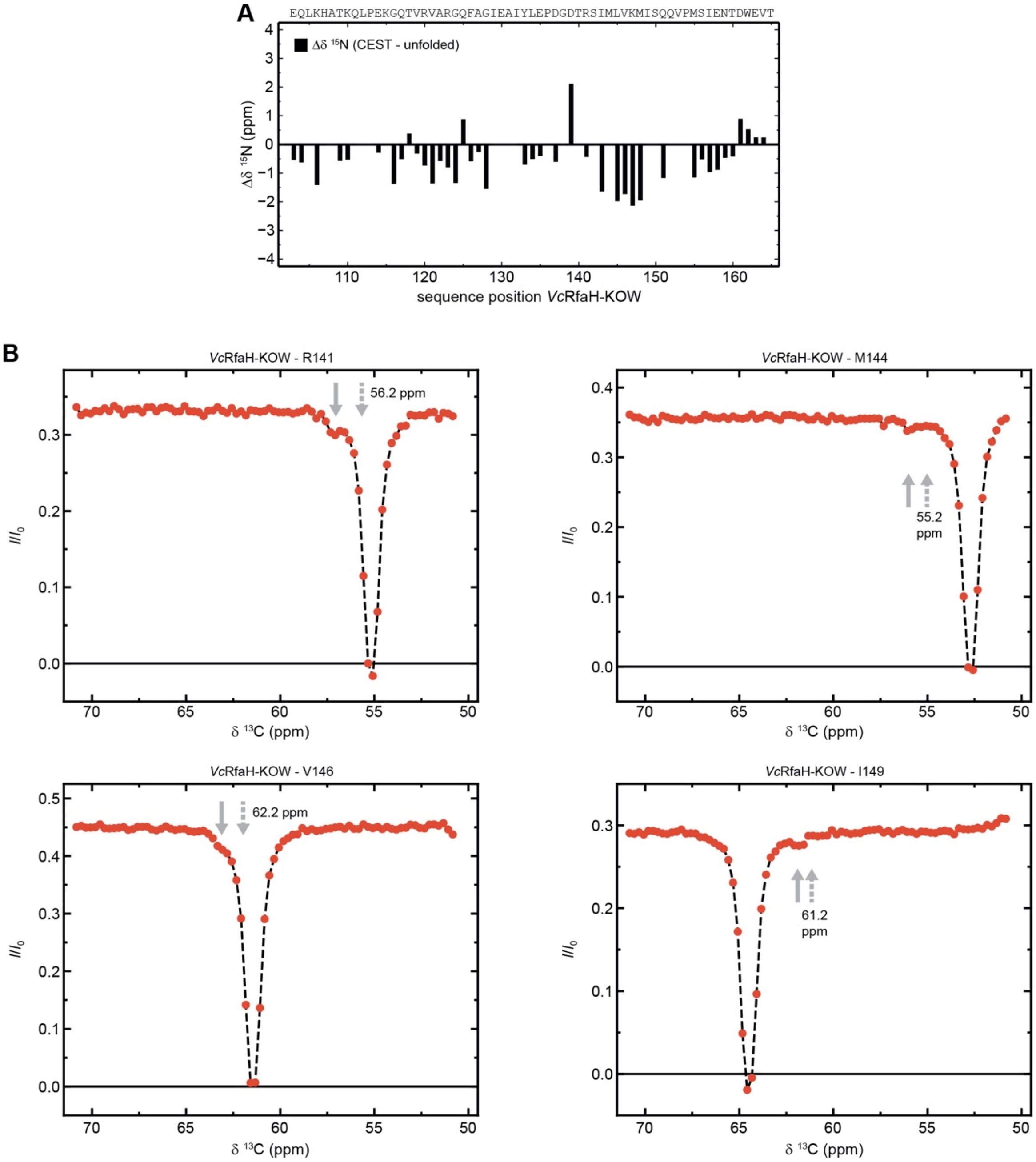


**Figure 6 – Figure supplement 3.** The minor species of *Vc*RfaH-KOW contains residual structure. (**A**) Sequence dependent difference between the $^{15}$N backbone amide chemical shifts of the CEST minor species of *Vc*RfaH-KOW and the corresponding theoretical random coil value. The sequence of the two protein construct is given above the diagrams. **(B)** Exemplary traces of

CEST experiments recorded on $^{13}C\alpha$ carbons of $^{13}C,^{15}N$-*Vc*RfaH-KOW. Solid arrows mark the positions of the minor species dips, dashed arrows indicate the predicted random coil values.

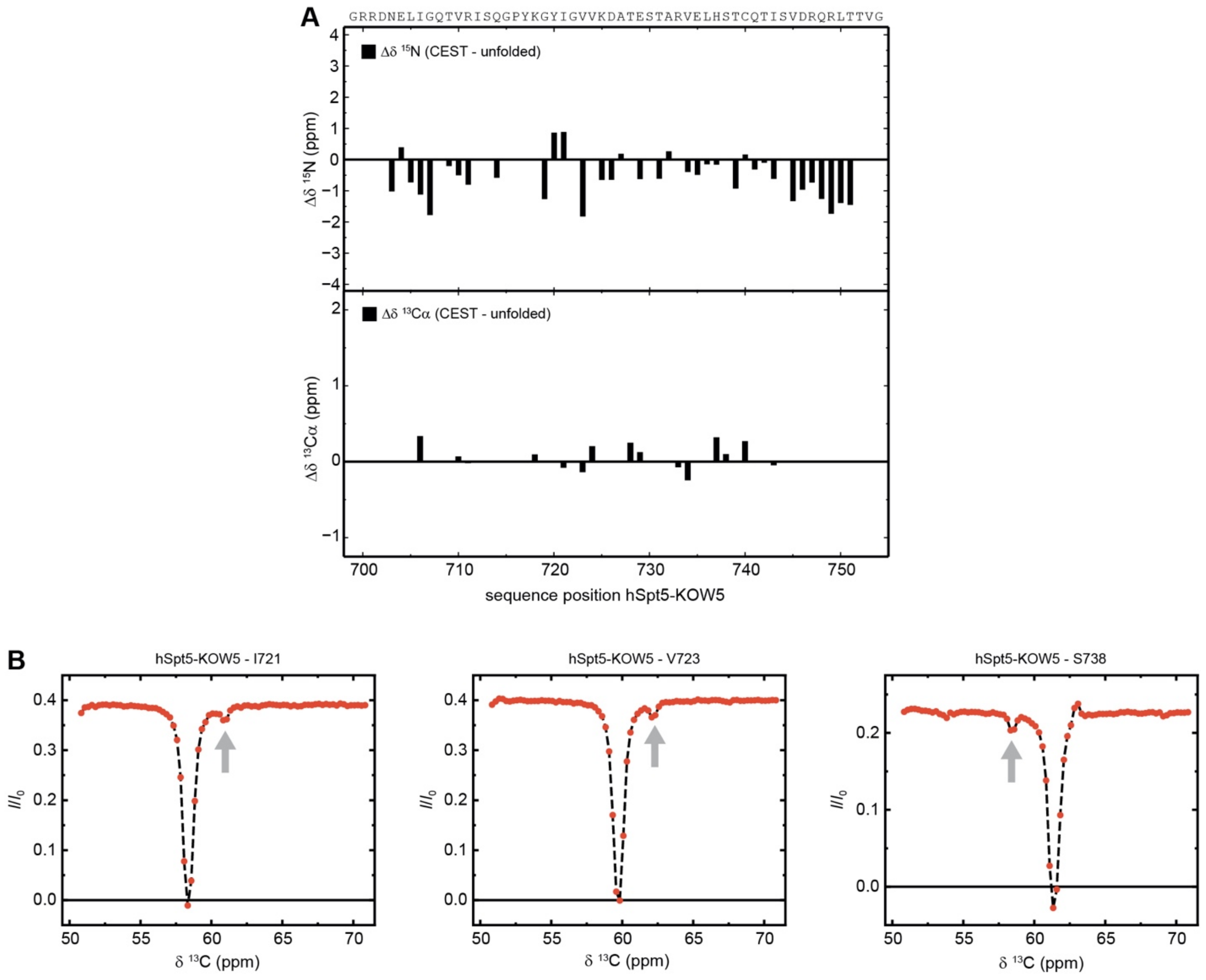


**Figure 6 – Figure supplement 4.** The minor species of hSpt5-KOW5 is completely unfolded. (**A**) Sequence dependent difference between the $^{15}N$ backbone amide (top) and $^{13}C\alpha$ carbons (bottom) chemical shifts of the CEST minor species of hSpt5-KOW5 and the corresponding theoretical random coil value. The sequence of the two protein construct is given above the diagrams. **(B)** Exemplary traces of CEST experiments recorded on $^{13}C\alpha$ carbons of $^{13}C$,$^{15}N$-hSpt5-KOW5. Arrows mark the positions of the minor species dips.

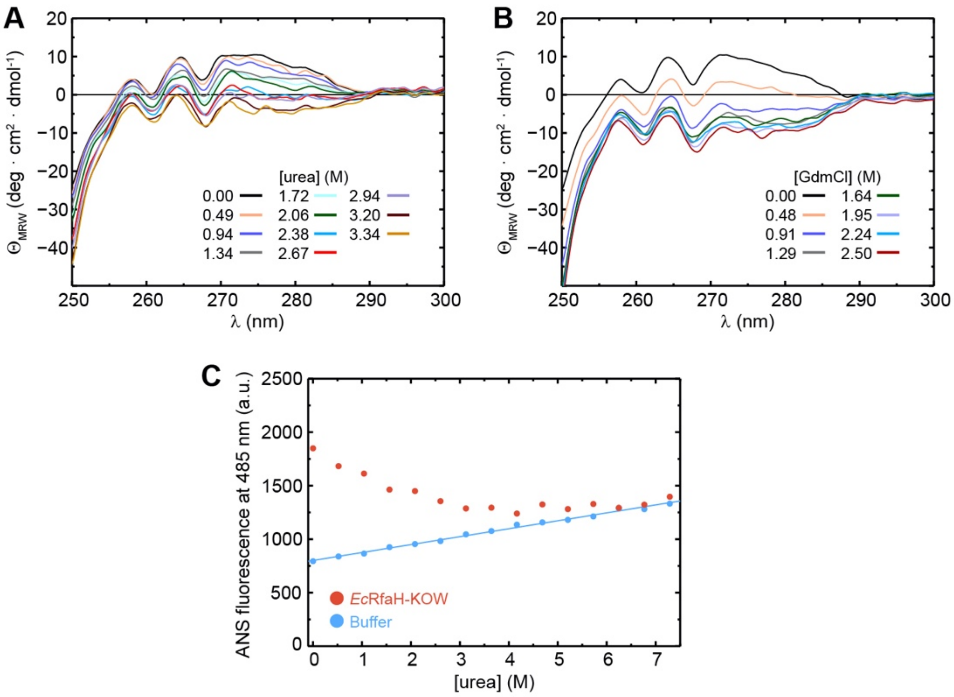


**Figure 6 – Figure supplement 5.** The intermediate state of *Ec*RfaH-KOW is no equilibrium MG. (**A, B**) Near-UV CD-spectra of *Ec*RfaH-KOW during a titration with (A) 10 M urea and (B) 8 M GdmCl. In both cases, the solution was buffered by 10 mM K-phosphate (pH 7.0). The denaturant concentrations at which the spectra were recorded are indicated. (**C**) ANS binding experiments. The graph shows the ANS fluorescence at 485 nm after over-night incubation of ANS in the presence (filled red circles) or absence (filled blue circles) of *Ec*RfaH-KOW at increasing urea concentrations. The system was buffered by 10 mM K-phosphate (pH 7.0).

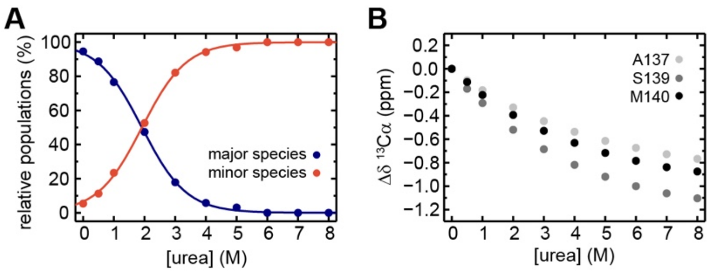


**Figure 6 – Figure supplement 6.** Extended analysis of the urea-induced denaturation of *Ec*RfaH-KOW. (**A**) Relative populations of the minor (filled red circles) and major (filled blue circles) species during the [$^{1}$H, $^{13}$C]-ctHSQC-based urea denaturation of $^{1}$H, $^{13}$C-*Ec*RfaH-KOW. The populations at a certain urea concentration were calculated from the ratio of volumes of the Hα/Cα correlation peaks of S139 minor or major species signals, respectively, to the sum of both values. The curves were fitted to a two-state model to extract the parameters of the transition from the major species to the minor species. The minor species was treated as a single species neglecting the fact that it is actually an ensemble of at least two subspecies. Fitting to a three-state (or even higher-state) model with an increased number of fitting parameters would not be appropriate due to the limited number of data points. (**B**) Chemical shift changes of $^{13}$Cα signals, Δδ$^{13}$Cα, of A137, S139 and M140 in the [$^{1}$H, $^{13}$C]-ctHSQC spectra during urea denaturation.
